# Genome Analysis of *Bacillus paralicheniformis* AA1 isolated from a conventional milpa farming system in the northwestern region of Sonora, Mexico

**DOI:** 10.1101/2025.01.08.631978

**Authors:** Andrés Chávez-Almanza, Eber Villa-Rodríguez, Jonathan Rojas-Padilla, Ernesto Cantú-Soto, Carlos Díaz-Quiroz, Abel Verdugo-Fuentes

## Abstract

This study provides an in-depth genome analysis of *Bacillus paralicheniformis* AA1, a bacterial strain isolated from a traditional milpa farming system in Sonora, Mexico. The genomic investigation revealed a high level of completeness, demonstrated by the presence of a diverse and functionally significant repertoire of genes associated with fundamental biological processes, including nutrient assimilation, stress response, and cellular regulation. Notably, the genome also contains genes responsible for the biosynthesis of secondary metabolites, highlighting its potential for biotechnological applications. Taxonomic classification was rigorously conducted using integrated genome-wide approaches, which definitively confirmed the identification of isolate AA1 as belonging to the species *Bacillus paralicheniformis*. Comparative genomic analysis further established a high degree of genetic similarity between AA1 and other *B. paralicheniformis* strains with well-characterized biotechnological capabilities. This similarity strongly suggests that AA1 harbors genetic elements responsible for the synthesis of antimicrobial compounds, enzymes with industrial relevance, and metabolites that promote plant growth. The findings underscore the potential of *Bacillus paralicheniformis* AA1 as a valuable resource for biotechnology and sustainable agriculture. By enhancing our understanding of microbial diversity within traditional agroecosystems, this study contributes to the broader knowledge base required for the development of innovative agricultural practices. Future research should focus on functional validation of key genes to fully unlock AA1’s potential as a bioresource for antimicrobial production, enzyme synthesis, and crop enhancement, paving the way for its application in environmentally sustainable farming systems.

## 1. Introduction

Bacteria exhibit a ubiquitous presence across diverse environments, spanning terrestrial and marine habitats, where they play diverse ecological roles (1). Among these, a range of different species within the genus *Bacillus* are commercially available biopesticides because of their capacity to synthesize a diverse group of antimicrobials, nematicidal, and insecticidal metabolites (2, 3)

In recent years, *Bacillus paralicheniformis* has gained attention for its remarkable characteristics, including its ability to produce antimicrobial bioactive substances and thrive in challenging environments (4, 5). From an agrobiotechnology perspective, *B. paralicheniformis* is an interesting source for biofertilizer development due to its multifaceted capabilities (6). These include the production of bioactive molecules that stimulate plant growth (*e*.*g*., indoleacetic acid, siderophores, ACC deaminase), enhancement of plant nutrition through nutrient solubilization (iron, phosphorus, and potassium), and inhibition of phytopathogen attacks either via antimicrobial production or by triggering plant immune responses (7, 8). This particular *Bacillus* species has demonstrated successful applications across a variety of economically significant crops, including tomato (9), mango (10), chili (11), highlighting its biotechnological relevance.

Milpa farming systems, characterized by the cultivation of multiple crop species such as maize, beans, herbs, and grass, have been linked to elevated soil microbial diversity in comparison to monocultures (12). In these highly diverse microbiomes, microbe-microbe interactions, involving both competition and cooperation, are naturally intensified. Consequently, microbes thriving in such environments often possess an interesting set of genes enabling them to effectively compete and collaborate with other microbial inhabitants, while also adapting to the selective pressures exerted by the varied root exudates of different plant species (13). This characteristic makes milpa production systems a promising reservoir of microbes with interesting biotechnological potential. In this work, we present the genome assembly of one of the most abundant bacterial isolates (strain AA1) recovered from a conventional milpa farming system in the northwestern region of Sonora, Mexico.

## 2. Materials and Methods

### 2.1 Sample collection

Soil samples were collected from a milpa production system at the Technological Institute of Sonora in the northwest region of Sonora, Mexico. The location is 27°29’46”N 109°58’18”W, and the elevation is 40 m above sea level. During May, the area typically receives temperatures of 27.5 °C and an average rainfall of 2.0 mm. During the vegetative phase of the maize growth cycle in May 2023, three composite soil samples weighing about 1 kg each were randomly obtained from three separate locations. The depth at which the samples were collected was 20 cm. The soil samples were subsequently examined for microbial analyses in the laboratory after being carried in a cooler maintained at 4 ± 2°C.

### 2.2 Microbiological analysis

*Bacillus* isolation and purification were carried out as follows; ten grams of each sample were dissolved in 90 mL of distilled water and heated to 80°C for 15 min. After diluting the solution with sterile water, 0.1 ml of the mixture was equally spread out on nutritional agar plates. The plates were then placed in an aerobic incubator at 37°C for 24 h. The colonies were then prepared for microscopic morphological identification using Gram staining and subjected to purification.

### 2.3 DNA extraction and sequencing

AA1 strain was grown in Trypticase soy broth (TSB) at 37 °C for 24 hours. Then, DNA was isolated using a commercial Blood and Tissue DNA kit (Qiagen Cat-69504) following the manufacturer’s guidelines. The purity and yield of DNA were spectrophotometrically determined utilizing Nanodrop 2000c (Thermo Fisher Scientific). The sequencing library was then constructed and sequenced using the Illumina NovaSeq platform. The experimental procedure was performed at The Sainsbury Laboratory in Norwich, United Kingdom, employing an axenic culture that had been previously supplied.

### 2.4 Bioinformatic analyses

Genome assembly was conducted using SPAdes genome assembler (version 3.15.4) (14). The resulting assembly was submitted to NCBI Prokaryotic Genome Annotation (PGAP) pipeline for i) prediction of protein-coding genes, ii) determination of RNAs, tRNAs, and pseudogenes, ii) gene annotation and iii) genome completeness and contamination assessment (15). Additionally, KOs and COGs assignments were executed using anvi’o v8 (16). Phylogenomics analysis was performed using GtoTree pipeline (17), while a pangenomics-based tree and ANI identity matrix were generated using anvi’o v8 (16). The genome map was generated using Proksee (18).

### 2.5 Data availability

The whole genome sequence was submitted to DDBJ/ENA/GenBank and assigned the accession number JAYMDM000000000. The sequenced strain’s BioProject database accession number is PRJNA1061142. The project’s Sequence Read Archive information can be accessed using the accession number SRR27458597.

## 3. Results and Discussion

Milpa farming systems are an interesting source of microorganisms with biotechnological potential. Therefore, we collected soil samples from a milpa production system and conducted bacterial isolation by serial dilution method. From this isolation effort, we identified strain AA1, one of the most abundant isolates present in the soil samples. We hypothesized that strain AA1 possesses a set of genes that enable it to thrive in this environment, which makes it an interesting target for whole genome analysis.

### 3.1 AA1 sequencing results in a high-quality genome assembly

To gain insights into the genetic features facilitating the proliferation of this strain within the studied milpa system, we isolated its genomic DNA (gDNA), sequenced it using the NovaSeq platform, and assembled the obtained sequencing reads using the SPAdes assembler. The novo genome assembly resulted in a total of 39 contigs, 10 contigs with sizes larger than 50 kb, and the largest contig of 1071 kb (Figure 1A and 1B). Next, we submitted our genome to the Prokaryotic Genome Annotation (PGAP) pipeline to identify genome features of AA1 (*e*.*g*., CDSs, RNAs, tRNAs, and pseudogenes) and perform genome completeness and contamination assessment. This analysis indicated high genome completeness (99.41 %) and no contamination (0%) (Figure 1C). Additionally, PGAP pipeline predicted i) a total of 4400 genes, of which 4212 correspond to protein-coding sequences and 75 pseudogenes, ii) 26 ribosomal RNAs (rRNAs), of which 9, 7, and 12 correspond to 5S, 16S, and 23S rRNA, respectively, and iii) 80 tRNAs and 5 ncRNAs (Figure 1C). According to NCBI Prokaryotic Genome Annotation Standards (19), the minimum standards to consider a genome “complete” are i) at least one copy of each rRNA (5S, 16S, 23S), ii) at least one copy of tRNAs for each amino acid and iii) Protein-coding genes count divided by genome length close to 1. Overall, our genome assembly surpasses the assemblies of previous studies (20, 21) and meets NCBI standards, indicating suitable assembly to explore the genomic information of this isolate.

**Figure 1.**
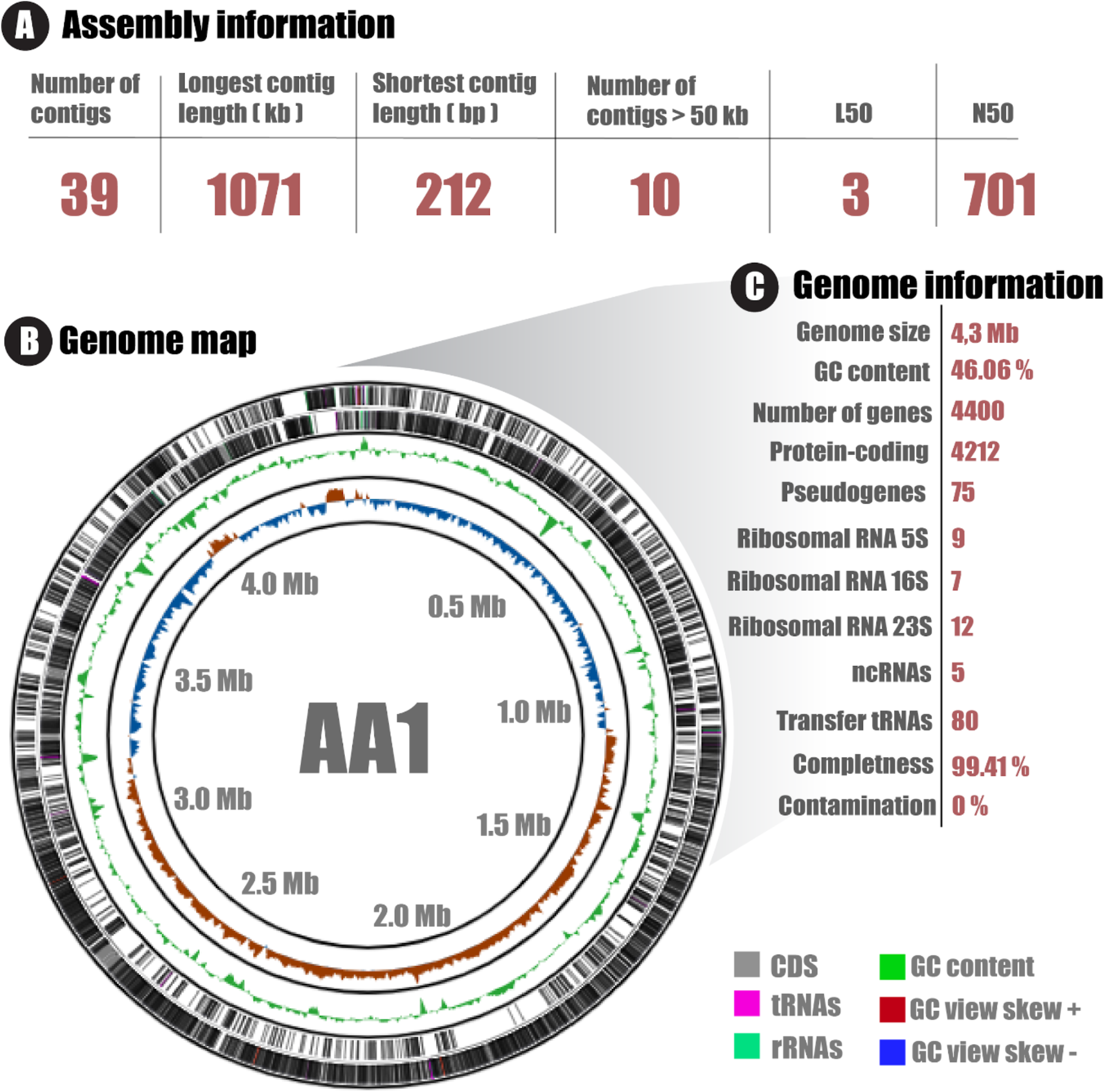
Genome assembly and statistics of AA1 genome. A) Quality information of AA1 genome assembly. B) AA1 circular genome map. From the innermost to the outermost rings; GC skew, GC content, CDS in minus (-) strand, CDS in plus (+) strand. C) AA1 genome information determined by PGAP pipeline.

### 3.2 Genome-wide analysis reveals AA1 is a *Bacillus paralicheniformis* strain

Taxonomic identification of bacterial isolates is a crucial step in bioprospection, as it enables us to gain insights into their biotechnological or pathogenic potential. In this work, we initially explore the taxonomic affiliation of strain AA1 by performing a BLAST using 16S rRNA, a highly conserved and widely used prokaryotic taxonomic marker (22). Thus, we observed that the 16S rRNA sequence of strain AA1 is highly conserved with some isolates of *B. licheniformis* (100 % identity/coverage) and *B. paralicheniformis* (100 % identity/coverage), suggesting that strain AA1 could belong to one of these taxonomic groups. While useful, 16S rRNA-based bacterial classification has the disadvantage that some bacteria may share high similarity with other members of the same family (23), as observed here.

Hence, we conducted an integrated genome-wide approach to taxonomically classify strain AA1 (Figure 2). First, we performed a phylogenomic analysis using 119 SCG markers commonly used in firmicutes (Figure 2A), alongside 38 *Bacillus* representative accessions retrieved from the Genome Taxonomy Database (24). This analysis facilitated the identification of different *Bacillus* species (*B. paralicheniformis, B. licheniformis, B. haynesii, B. sweneyi, B. pumilus*) closely related to AA1 strain. Subsequently, a pangenome-based tree was constructed using high-quality complete genomes (including type strains) of closely related species, obtained from EZbiocloud (www.ezbiocloud.net). This approach clusters genomes based on the presence or absence of predicted coding regions. Our pangenome-based tree reveals three distinct groups, with the AA1 genome present in group 1, clustering alongside *B. paralicheniformis* genomes. This clustering strongly suggests AA1 affiliation with this taxonomic group. Lastly, we conducted an Average Nucleotide Identity (ANI) analysis using genomes within group 1. In bacterial taxonomy, species are typically distinguished based on an ANI threshold of 95% (25). Our analysis revealed that AA1 exhibits an ANI value higher than 95% when compared to any of the *B. paralicheniformis* genomes included in the study. This result shows that strain AA1 can unequivocally be classified as *Bacillus paralicheniformis*. Noteworthy, this *Bacillus* species has been associated with multiple traits with biotechnological relevance (8, 10), suggesting that strain AA1 could harbor some of these traits.

**Figure 2.**
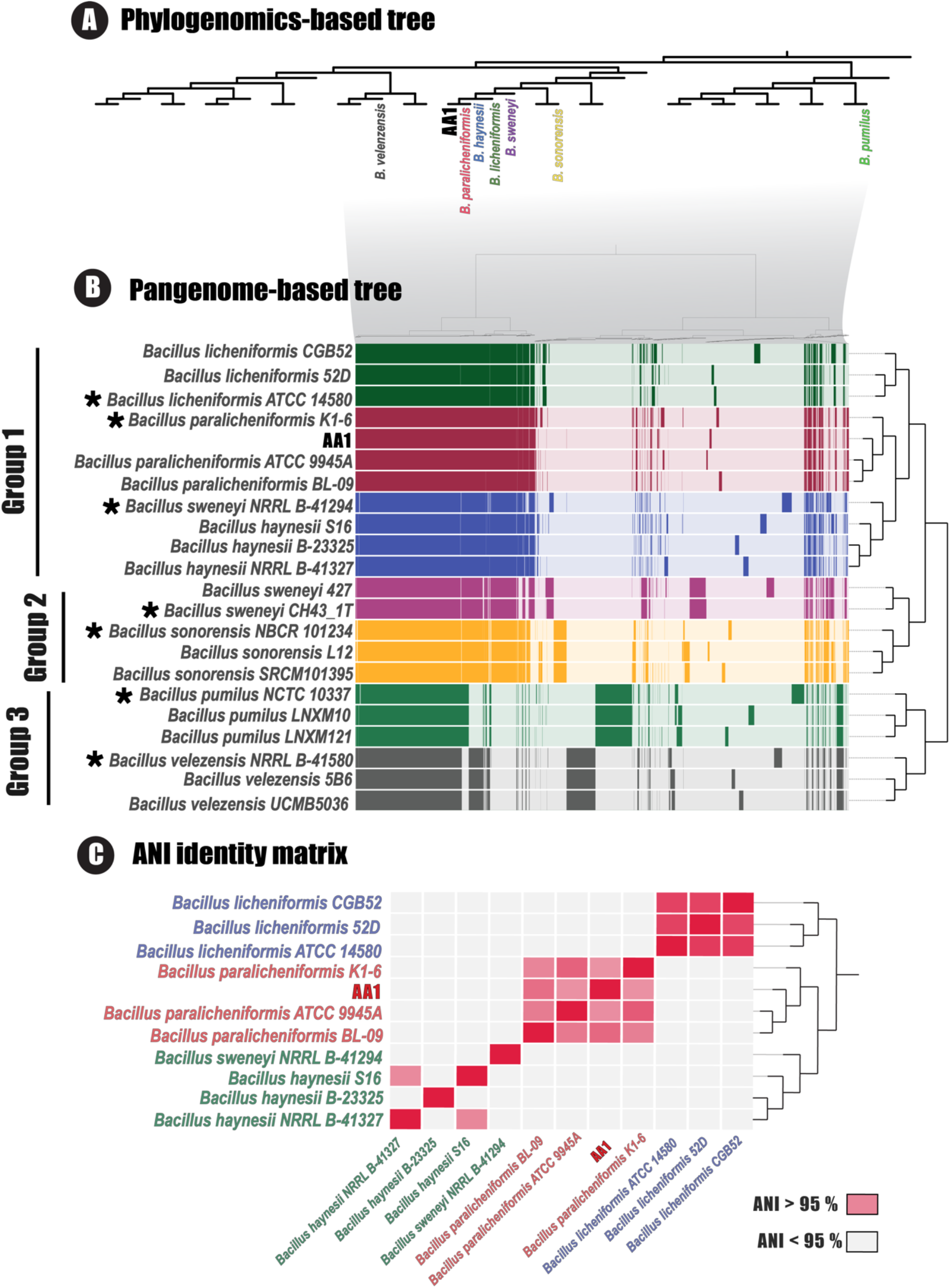
Genome-wide analysis determines AA1 taxonomy. A) Phylogenomic analysis using a single copy gene (SCG) set for Firmicutes (119 genes). B) Pangenome-based tree using closely related species determined in phylogenomic tree using SCG. Asterisks denote type strains retrieved from EZbiocloud (www.ezbiocloud.net). Each line represents a genome. Filled-colored lines (Green to gray for *B. licheniformis, B. paralicheniformis, B. haynesii, B. sweneyi, B. pumilus*, and *B. velezensis*, respectively) represent the presence of a gene, while lighter color represents the absence of a gene. C) ANI identity matrix using strains present AA1 clade in the pan genome-based tree.

### 3.3. Genome annotation and comparative genomics suggest *Bacillus paralicheniformis*

#### AA1 has biotechnological potential

In bacterial bioprospection, genome annotation is an essential step since it facilitates the identification of the functional elements found in the genome. This process provides valuable insights that could reveal potential biotechnological applications for bacterial isolates. In this sense, we perform the genome annotation of *Bacillus paralicheniformis* AA1 to gain insight into their potential biological functions. To this end, we use an integrated approach using different annotation systems (PGAP, KEGG, COG). A total of 3827, 3427, and 2971 genes were annotated with PGAP, COG, and KEGG systems, respectively, displaying multiple overlaps in the annotated genes (Figure 1A). Overall, with our approach, we were able to retrieve biological information from around 91 % of the coding sequences (Figure 1A).

Within the AA1 genome, there is an overrepresentation of genes linked to fundamental biological processes essential for cellular function. These include genes involved in i) carbohydrate and metabolism (345 genes), ii) transcription (305 genes), iii) amino acid metabolism and transport (303 genes), and iv) translation, ribosomal structure, and biogenesis (245 genes). Noteworthy, *Bacillus paralicheniformis* AA1 has 81 genes involved in secondary metabolite biosynthesis (Figure 3B), which suggests that this strain could produce several metabolites with biotechnological potential.

**Figure 3.**
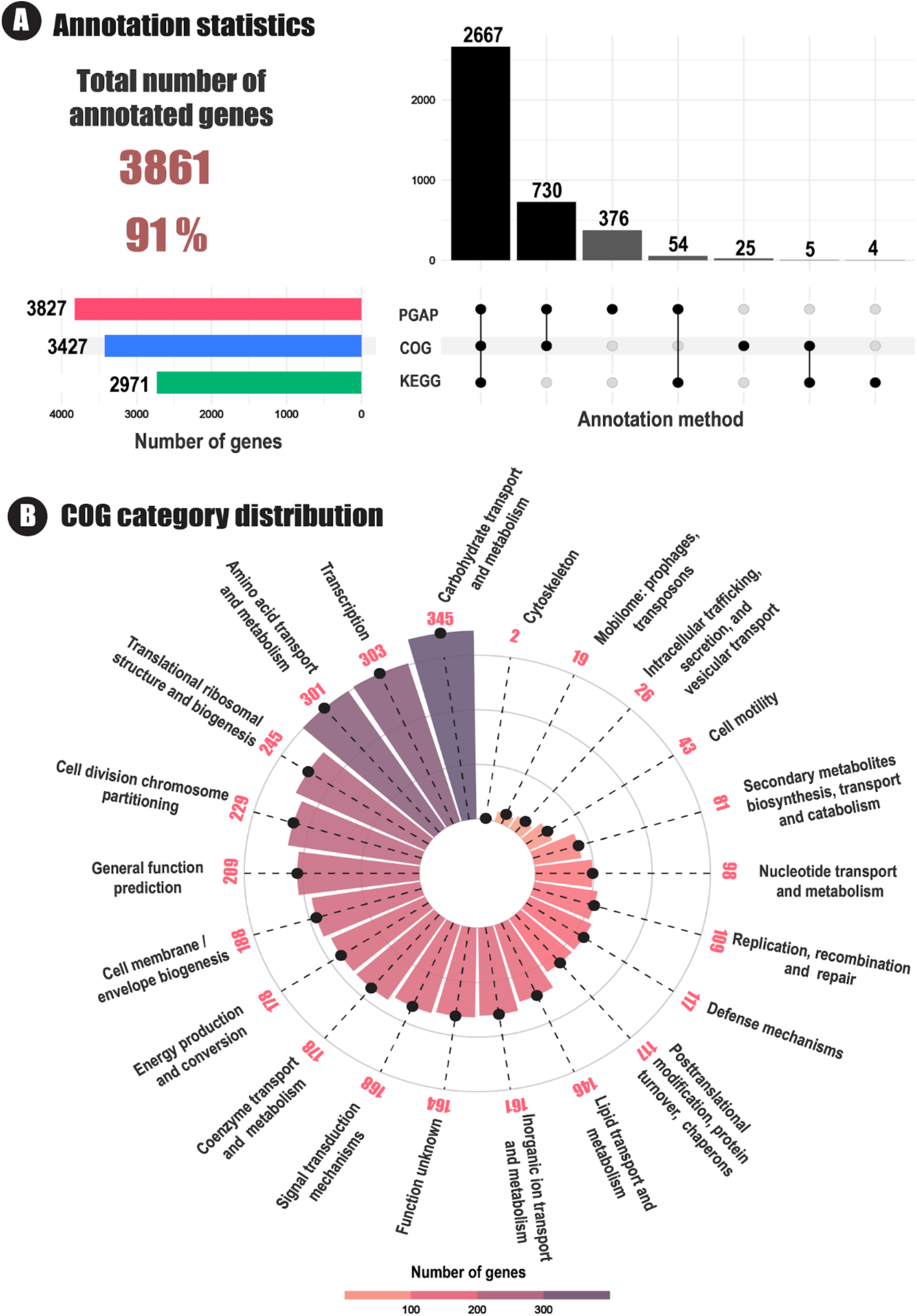
Genome annotation of *Bacillus paralicheniformis* AA1. A) Upset plot displaying overlaps of gene annotations by multiple annotation systems. B) Distribution of Clusters of Orthologous Genes (COG) categories within the AA1 genome, with respective gene counts indicated.

In the last years, several *Bacillus paralicheniformis* representatives with biotechnological potential have been studied. This can be exemplified in a recent study where a comprehensive genome analysis of *Bacillus paralicheniformis* BP9 revealed this strain has multiple interesting biotechnological traits, such as production of secondary metabolites (bacillibactin, fengycin, bacitracin, and lantibiotics), genes involved in niche competition, genes involved in plant growth promotion (26). Similarly, genome-wide analysis in other *Bacillus paralicheniformis* strains, TRQ65 and KMS 80, evidenced similar traits (27, 28). In this work, we perform a comparative genomic analysis using AA1 alongside these strains exhibiting biotechnological traits (Figure 4). Our analysis reveals that while each strain harbors a unique set of genes, there exists a high overlap (ANI > 99 %), with more than 4300 genes shared among them. Furthermore, we observe that strain AA1 shares a closer genomic content, particularly in terms of gene presence/absence, with the BP9 strain. The high overlap of genes between strain AA1 and other *Bacillus paralicheniformis* strains known for their biotechnological applications indicates that AA1 harbors promising genetic elements. Nonetheless, a future analysis focusing on identifying and validating AA1 biotechnological traits must be undertaken.

**Figure 4.**
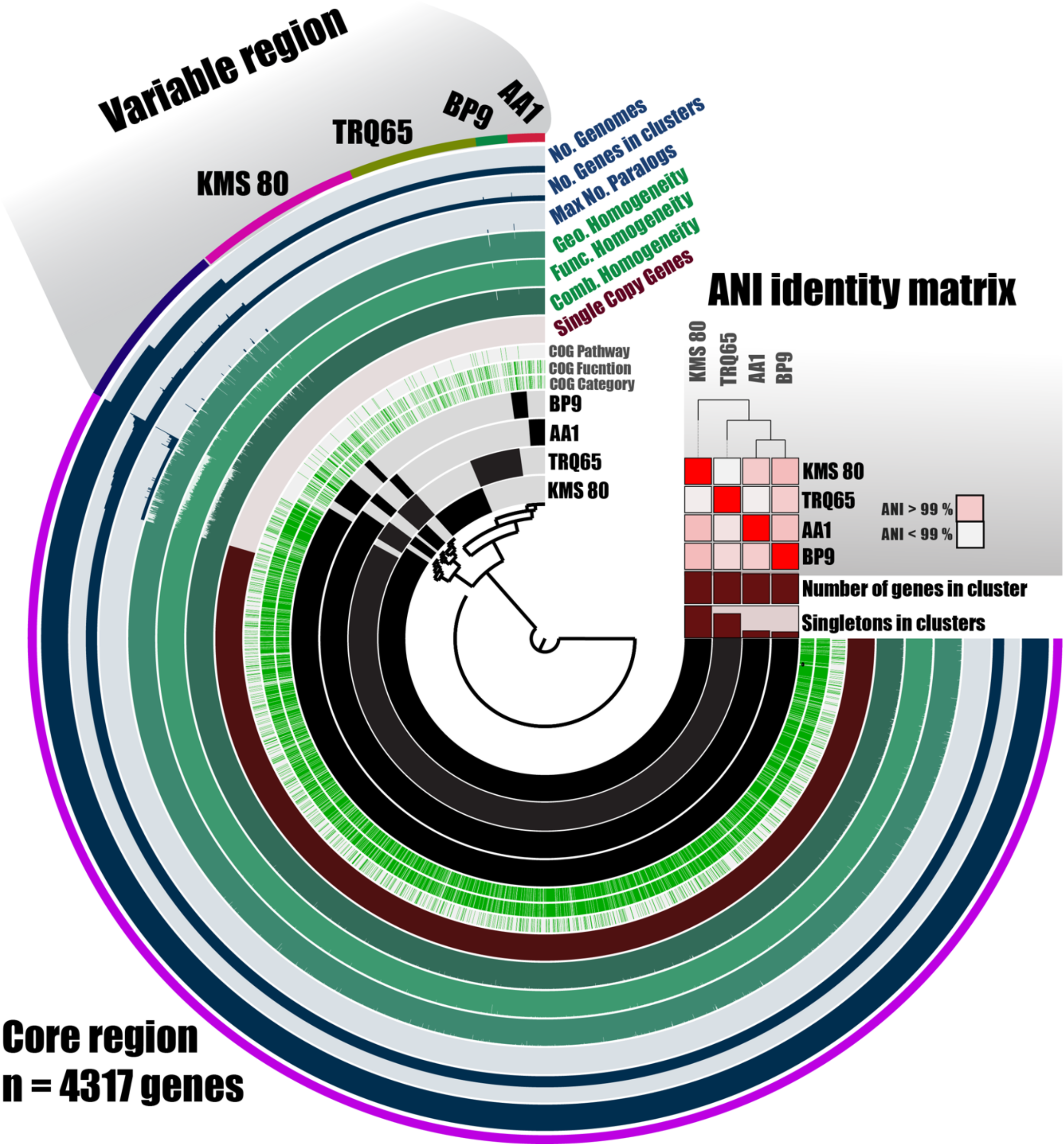
illustrates the comparative genomics of Bacillus paralicheniformis AA1 alongside other strains known for their biotechnologically relevant traits within the species. Each line in the graph represents a genome, with black-filled segments indicating the presence of a gene and lighter sections denoting gene absence. Additionally, unique genes present in the analyzed genomes are specifically highlighted.

## Conclusion

The genome analysis of *Bacillus paralicheniformis* AA1, isolated from a milpa farming system in Sonora, Mexico, reveals a wealth of genetic features that underscore its potential biotechnological applications. This study enhances our understanding of the microbial diversity associated with milpa ecosystems and highlights the capacity of such microorganisms to contribute to sustainable agricultural practices. The findings lay a foundation for exploring AA1’s functional roles, particularly in antimicrobial compound synthesis, industrial enzyme production, and plant growth promotion. Future research focused on identifying and characterizing specific genes responsible for these traits will be pivotal in fully realizing the biotechnological promise of *Bacillus paralicheniformis* AA1, potentially offering innovative solutions for agriculture and biotechnology.

## Funding information

The sequencing of the bacterial strains was supported by GetGenome and The Sainsbury Laboratory, Norwich, UK, with support from the Gatsby Charitable Foundation and the Biotechnology and Biological Sciences Research Council (BBSRC).

## Conflict of Interest

The authors declare that there are no conflicts of interest.

---

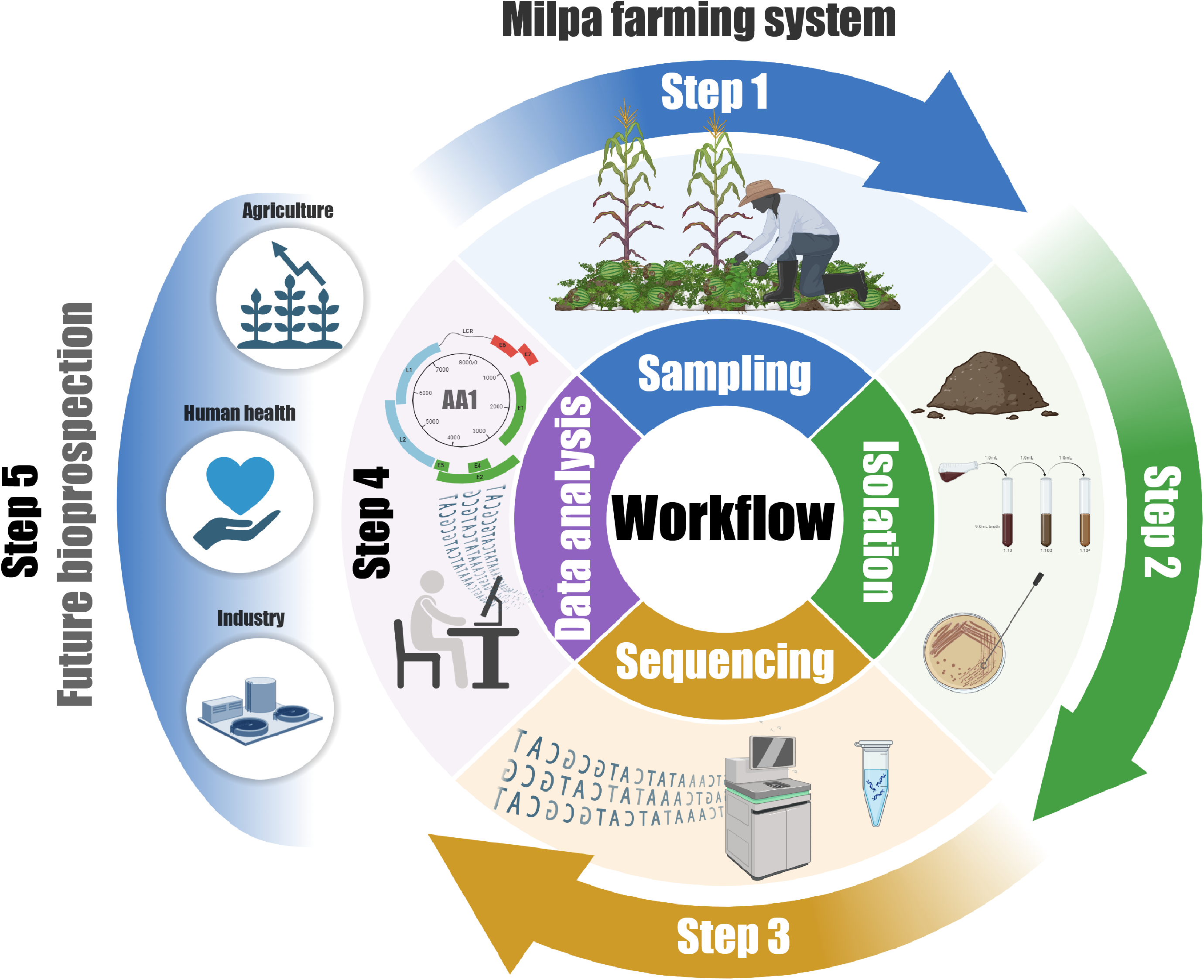

## Notes

### Competing Interest Statement

The authors have declared no competing interest.

